# Inferring diploid 3D chromatin structures from Hi-C data

**DOI:** 10.1101/644294

**Authors:** Alexandra Gesine Cauer, Gürkan Yardimci, Jean-Philippe Vert, Nelle Varoquaux, William Stafford Noble

## Abstract

The 3D organization of the genome plays a key role in many cellular processes, such as gene regulation, differentiation, and replication. Assays like Hi-C measure DNA-DNA contacts in a high-throughput fashion, and inferring accurate 3D models of chromosomes can yield insights hidden in the raw data. For example, structural inference can account for noise in the data, disambiguate the distinct structures of homologous chromosomes, orient genomic regions relative to nuclear landmarks, and serve as a framework for integrating other data types. Although many methods exist to infer the 3D structure of haploid genomes, inferring a diploid structure from Hi-C data is still an open problem. Indeed, the diploid case is very challenging, because Hi-C data typically does not distinguish between homologous chromosomes. We propose a method to infer 3D diploid genomes from Hi-C data. We demonstrate the accuarcy of the method on simulated data, and we also use the method to infer 3D structures for mouse chromosome X, confirming that the active homolog exhibits a bipartite structure, whereas the active homolog does not.

## 1 Introduction

The 3D organization of the genome plays an important role in regulating basic cellular functions, including gene regulation [29, 33], differentiation [20, 11], and the cell cycle [26]. Chromosome conformation capture techniques such as Hi-C measure the frequency of interactions between pairs of loci, thereby allowing a systematic analysis of genome structure. Although Hi-C contact matrices yield valuable insights, modeling and visualizing genome structures in 3D can unveil relationships and higher-order structural patterns that are not apparent in the raw data [12, 36, 26, 23] by providing a humanly interpretable 3D structure, orienting genomic regions relative to various nuclear landmarks, and serving as a framework for integrating other data types [5]. Embedding contact count data in a 3D Euclidean space can also reduce noise in the underlying Hi-C data.

Previous methods to inferring chromatin structure from population Hi-C data fall into one of two broad categories. “Ensemble” approaches create populations of 3D structures that jointly explain the observed Hi-C data [28, 36, 7, 16, 19, 35, 14, 24, 38, 39, 18]. Theoretically, structural ensembles can mimic the heterogeneity of cells in a population. However, these methods are frequently underdetermined because there are often more parameters to estimate for a large population of cells than data points. Ensemble models can also be difficult to validate and interpret. “Consensus” approaches, on the other hand, make the assumption that bulk Hi-C data can be accurately summarized in a single, consensus 3D structure [12, 9, 34, 37, 2, 22, 15]. Modeling a single structure tends to be less computationally demanding than modeling an entire population of structures. Furthermore, the resulting model has the advantage of relatively straightforward visualization and interpretation.

For either ensemble or consensus approaches, a particular challenge is presented by Hi-C data derived from diploid organisms. As in most high-throughput sequencing experiments, a typical Hi-C experiment does not produced phased data; that is, the data does not distinguish between allelic copies. Thus, an observation of a single Hi-C contact between loci *i* and *j* corresponds to one of four possible events: either copy of locus *i* coming into contact with either copy of locus *j*. Any 3D inference method that aims to model diploid genomes must accurately account for this allelic uncertainty.

A variety of strategies have been developed to account for diploidy in Hi-C 3D models. In general, ensemble models face less of a challenge on this front, since the two allelic copies can be treated like additional members of the ensemble. Among consensus methods, by far the most common approach is to assume that the two homologous copies of a given chromosome share the same 3D structure [37, 22, 40] and then to model each chromosome separately.

We are aware of only three previous attempts to model diploidy in non-ensemble methods. Previously, we described an extension of our PASTIS software to handle the near-haploid cell line KBM7 [3]. We proposed to infer jointly the distribution of contact counts between homologs and the 3D structures by maximizing a constrained and relaxed likelihood. However, this relaxation is unsatisfying, as it yields non-integer counts modeled as random Poisson variables. More recently, two separate research groups have developed methods for modeling diploid genomes from single-cell data [6, 32]. However, these methods cannot be directly applied to bulk Hi-C data, which is much more widely available.

In this work, we propose a method to infer diploid consensus 3D models from Hi-C data. Our approach builds upon PASTIS [37], which infers 3D models by using a Poisson model of Hi-C counts coupled with a simple biophysical model of polymer packing. The key idea of extending PASTIS to infer diploid genomes is to explicitly model the uncertainty of allelic assignments for each observed read. We consider two distinct settings: the more challenging setting where the data is fully ambiguous, and the setting where a subset of the reads can be mapped to a single parental allele. To assist in inference, we incorporate several constraints into our objective function, reflecting our prior knowledge of genome architecture. Through extensive simulations, we demonstrate that our approach can successfully model two distinct homologous chromosome structures, given a sufficient number of reads, even when the data is fully ambiguous. We also apply our approach to real Hi-C data derived from a first generation (F1) cross of two divergent mouse strains (F121 and *Castaneus*). The resulting diploid model of the X chromosome exhibits the expected “superdomain” structure [10], and is quite distinct from the inferred structure of the inactive X.

## 2 Method

Hi-C experiments involve sequencing pairs of interacting DNA fragments. Specifically, cells are cross-linked, DNA is digested using a restriction enzyme, and interacting fragments are then ligated together. Fragments are subsequently sequenced through paired-end sequencing, and each mate is associated with one interacting locus. Hi-C data can then be summarized in a symmetric *n* × *n* contact count matrix *C*, where each row and column corresponds to a genomic locus and each matrix entrix *c*_*ij*_ to the number of time those two loci have been observed to interact.

For diploid organisms, reads from homologous chromosomes cannot be distinguished from one another, and the resulting Hi-C matrix aggregates contact counts from homologous chromosomes into a single Hi-C matrix (Figure 1). The challenge of inferring diploid structures from Hi-C data lies in disambiguating the contact counts from the two homolog chromosomes. We call these aggregated counts “ambiguous,” and denote by *C*^*A*^ the corresponding contact count matrix. If the parental genomes are known *a priori*, then a small proportion of reads can be mapped to each haplotype: contact counts from the two homolog chromosomes can be disambiguated based on heterozygous positions, yielding a single-allele Hi-C count matrix [29, 10]. We refer to these counts as “unambiguous” and denote the corresponding matrix by *C*^*U*^. On the other hand, if only one mate can be mapped uniquely to one of the homologous chromsosome, then the contact count is only partially disambiguated between the two homologs. We refer to these as “partially ambiguous” contact counts, and we denote the corresponding matrix by *C*^*P*^.

**Figure 1:**
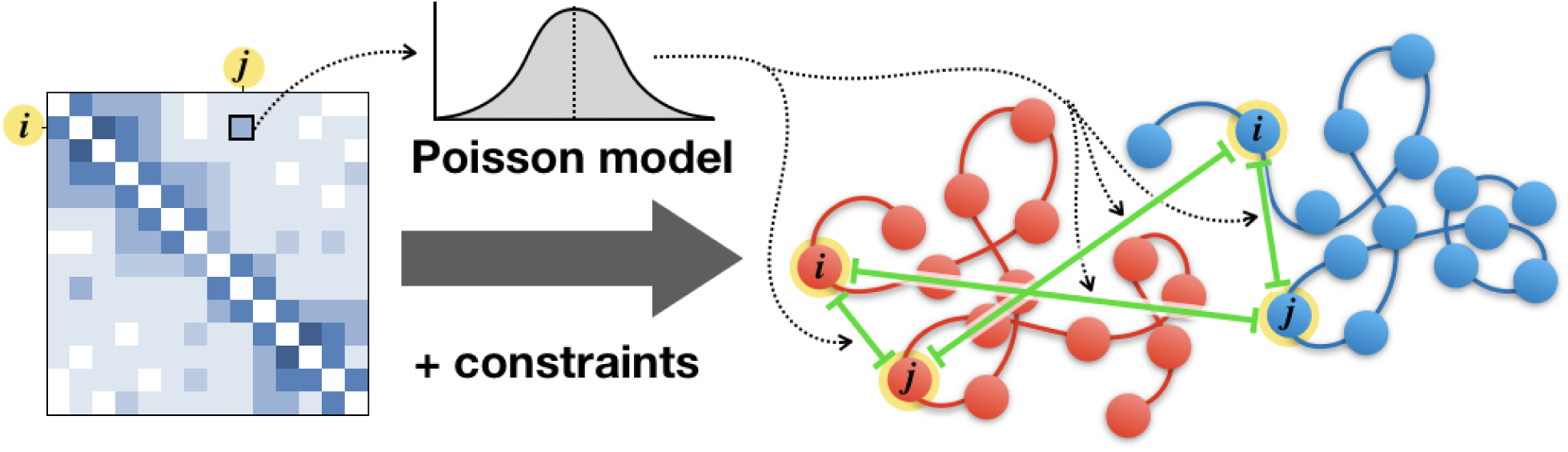
Inferring 3D structure using ambiguous diploid data. Each observed count (left) corresponds to a sum of four pairs of genomic loci (right). The Poisson model must be adjusted to account for this ambiguity.

We model chromosomes as *m* evenly-spaced beads, and we denote by **X** = (*x*_1_,…,*x*_*m*_) ∈ ℝ^3×*m*^ the coordinate matrix of the structure. The variable *m* denotes the total number of beads in the genome, and *x*_*ℓ*_ ∈ ℝ^3^ corresponds to the 3D coordinate of the *ℓ*-th bead. In the case of a haploid structure, the number of beads corresponds to the number of rows and columns in the contact count matrix *C*: *n* = *m*.

### 2.1 Inferring haploid structures with a Poisson model

Before we turn to inferring diploid structures, let us first review the approach proposed by PASTIS [37] to infer haploid 3D structures from a bulk Hi-C contact map **C**. PASTIS models the interaction frequency between genomic loci *i* and *j* as a random independent Poisson variable, where the intensity of the Poisson distribution is a decreasing function *f* of the distance between the two beads. Leveraging relationships found from studying biophysical properties of DNA as a polymer, PASTIS sets this function as follows: 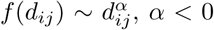. The *α* parameter can be set using prior knowledge (e.g., *α* = −3), or inferred jointly with the 3D structure. Inference is thus performed by maximizing the likelihood of the following Poisson model:

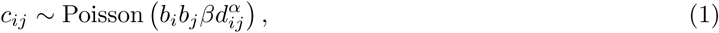

where *β* scales for the total number of contacts in the matrix (“coverage”), and *b*_*i*_ and *b*_*j*_ are locus-specific biases that are estimated using a standard procedure [17].

Our strategy to infer diploid structures builds upon this approach. Note that inferring a diploid structure from “unambiguous” contact counts *C*^*U*^ is similar to inferring a haploid structure from a classic Hi-C experiment, with the only difference concerning the biases, which are computed using all contact counts available per locus.

### 2.2 Modeling contact counts of diploid structures with a Poisson model

We propose to extend PASTIS to diploid genomes by leveraging the properties of each type of Hi-C contact map: ambiguous, partially ambiguous, and unambiguous. Let us first take a closer look at the common scenario, where the data is fully ambiguous.

For a given ambiguous contact count matrix *C*^*A*^, each observed contact count 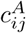 between a given pair of loci (*i, j*) corresponds to the sum of four different unambiguous contact counts (Figure 1):

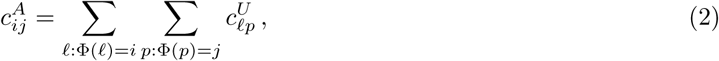

where Φ: [1, *n*] → [1, *m*] is the mapping that associates bead *ℓ* with locus *i*. Leveraging the property that the sum of *i* Poisson variables of intensities *λ*_*i*_ is a Poisson variable of intensity Σ_*i*_ *λ*_*i*_, we model the interaction count as

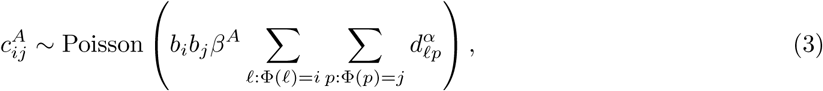

where *m* is the number of loci, *n* is the number of beads, *d*_*ℓ,p*_ is the Euclidean distance between beads *ℓ* and *p*, and *β*^*A*^ is a scaling factor determined by the coverage of the ambiguous contact count matrix.

Similarly, for a given partially ambiguous contact count matrix *C*^*P*^, each observed contact count 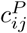 between a given pair of loci corresponds to the sum of two unambiguous contact counts, and is modeled by the interaction frequency of two pairs of loci.

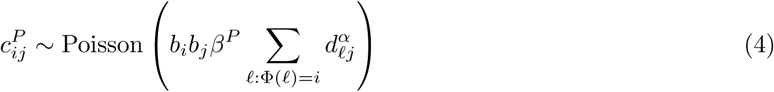

*β*^*P*^ is a scaling factor determined by coverage of the partially ambiguous contact count matrix.

We can thus cast the 3D structure inference as maximizing the log-likelihood

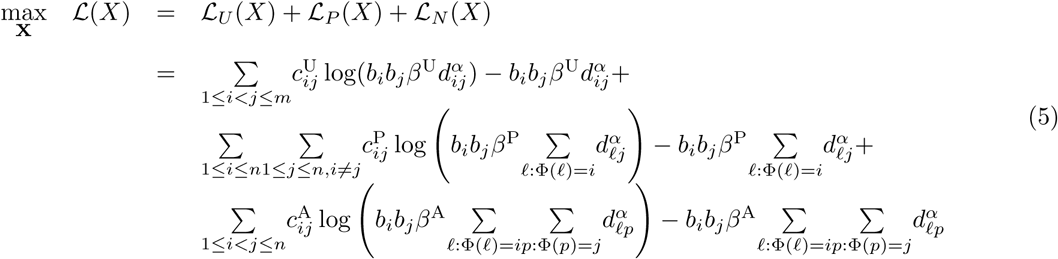

Note that this approach holds for polyploid genomes in addition to diploid genomes.

### 2.3 Incorporating prior knowledge

Because the resulting optimization is challenging, we add two constraints that reflect our prior knowledge about chromatin 3D structure: two neighboring beads should not be too far apart from one another, and homologs of most organisms occupy distinct territories [32, 31, 4, 27].

The first constraint maintains bead chain connectivity by minimizing the variance in the distance between neighboring beads:

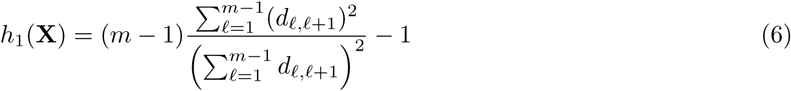

where *ℓ* and *ℓ* + 1 are on the same chromosome. This type of constraint has been used previously in Simba3D [30].

The second constraint aims to disentangle the structures of the two homologs, and operates on the distance between homolog centers of mass:

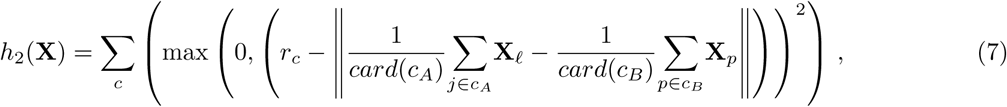

where *c* denotes the chromosome, *c*_*A*_ and *c*_*B*_ denote the set of beads associated to the two homologs of chromosome *c*, and *r*_*c*_ is a predefined scalar that increases relative to the space between the two homologs of chromosome *c*. We note that such a penalty may be interpreted as a log-prior in a Bayesian setting, where the distance between homolog centers of mass of chromosome *c* is *a priori* normally distributed with mean *r*_*c*_.

When unambiguous data is available, the values of *r*_*c*_ may be estimated via the distances between homolog centers of mass in an extremely coarse-grained structure inferred from unambiguous data alone. Alternatively, when unambiguous data is not available, *r*_*c*_ may be estimated as the mean distance between chromosome centers of mass in a coarse-grained structure inferred from ambiguous data, since this distance is expected to be similar to that between homologs.

We penalize the likelihood in Equation 5 and solve the following optimization problem:

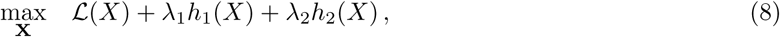

where *λ*_1_ and *λ*_2_ are penalization parameters, the values of which were chosen via a grid search. A version of PASTIS that implements the diploid inference approach is available at https://github.com/hiclib/ pastis.

### 2.4 Data

#### 2.4.1 Simulated Hi-C data

To validate our approach, we generated 10 simulated genomes with coverage, number of beads, and ratios of disambiguated contact counts corresponding to those of Hi-C data from the mouse Patski cell line (described in Section 2.4.2) at 500kb resolution. We also generated additional sets of 10 simulated genomes with the same number of beads, varying the proportion of ambiguous, unambiguous, and partially ambiguous contact counts.

To simulate “true” structures, we applied a random walk algorithm. This algorithm places beads successively along each chromosome, constraining each bead to lie within a given distance of the previous bead, provided the new bead does not overlap with any of the previously placed beads and that the entire homolog fits within a sphere of a predefined radius. We then derive unambiguous counts using the following model:

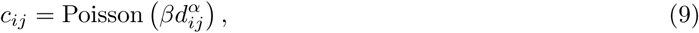

where *α* = −3, corresponding to a previously used theoretical exponent for the contact-to-distance transfer function [37]. Subsequently, we converted a portion of unambiguous counts to ambiguous or partially ambiguous counts by summing contacts from the appropriate pairs of loci.

#### 2.4.2 Real Hi-C data

We applied our method to publicly available in situ DNAse Hi-C of Patski fibroblast mouse kidney cells [10]. This line was derived from F1 female embryos, obtained by mating a BL6 female with a *Spretus* male. The BL6 female had an *Hprt* mutation, so hypoxanthine-aminopterin-thymidine medium was used to select for cells with X chromosome inactivation on the maternal allele.

### 2.5 Structure similarity measures

We use the following quantitative measures of similarity between 3D structures to determine the quality of structures inferred from simulated data and assess the stability of chromatin structures across biological replicates.

Root mean square deviation (RMSD) is a common way of comparing two three dimensional structures described by their coordinates **X, X**′ ∈ *R*^3×*m*^. RMSD is defined as

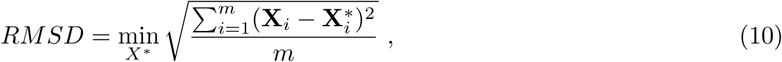

where **X**\* is obtained by translating, rotating, and rescaling **X**′ (**X**\* = *s***RX**′−**t** where **R**∈*R*^3×3^ is a rotation matrix, **t**∈*R*^3^ is a translation vector, and *s* is a scaling factor). RMSD values are computed independently on each homolog of each chromosome and summed.

Distance error [37] assesses the similarity between two distance matrices. This measure assigns more weight to long distances than RMSD. It is given by

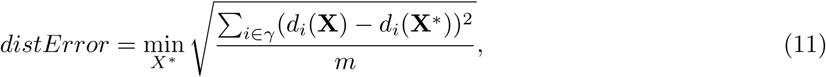

where *γ* is a set of distances of interest (e.g., intra-chromosomal distances). The structure **X**\* is obtained by rescaling **X**′(**X**\* = *s***X**′ where *s* is a scaling factor). To distinguish discrepancies in intra-chromosomal structure from those affecting the relative orientation of each pair of homologs or the relative orientation of different chromosome pairs, we compute distance error in two ways. Intra-chromosomal distance error is computed separately for each homolog of each chromosome, and *γ* encompasses distances between all beads of the given homolog. Inter-homolog distance error is computed separately for chromosome pair, and *γ* encompasses distances connecting all beads of two different homologs of a given chromosome. For both measures, values are summed for all chromosomes.

## 3 Results

### 3.1 Constraints improve ambiguous inference

First, we assessed the accuracy of our method on simulated datasets (Section 2.4.1) using ambiguous data alone, with and without our proposed constraints. Because of the lack of disambiguated contact counts, we expected this inference task to be difficult. Our results demonstrated that the two sets of constraints—bead connectivity and homolog separation—are necessary for successful inference. In the absence of the constraints, ambiguous inference performed poorly (Figure 2). Specifically, inferred homolog structures overlapped one another, and adjacent beads sometimes had large gaps between one another. The homolog separation constraint (Equation 7) and the bead connectivity constraint (Equation 6) were specifically designed to address these problems. Therefore, we repeated the inference with each constraint individually and the two constraints in combination. In this experiment, we compared results generated with and without each constraint at the optimal *λ* values (*λ*_1_ = 10^8^ and *λ*_2_ = 10^6^, respectively). The results showed that RMSD and distance error are lowest when both constraints were incorporated (Figure 2), and error scores obtained from structures inferred with both constraints were significantly lower than those obtained from structures inferred without constraints (pairwise t-test, Bonferroni corrected *p*-value <0.05, Supplementary Table S1). 3D structures produced with the constraints had fewer large gaps between neighboring beads and exhibited distinct territories for the two homologs.

**Figure 2:**
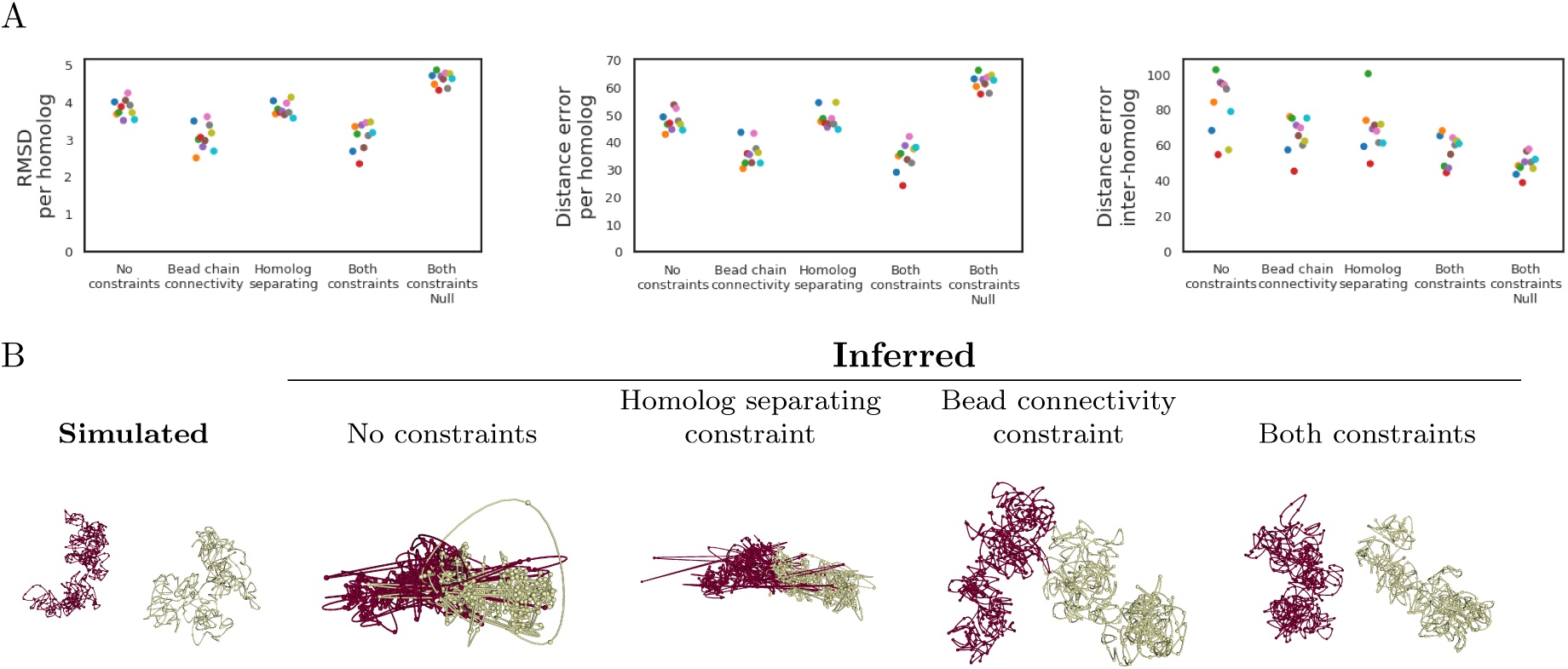
Constraints improve ambiguous inference. The simulated data consists of a single diploid chromosome with 9.3×10^6^ reads and 343 beads, the size of which corresponds to mouse chromosome X at 500 kb resolution. (A) The quality of the inferred structure, as measured by three different error scores (y-axis), improves upon application of the bead connectivity constraint (*λ*_1_ = 10^8^) and the homolog separating constraint (*λ*_1_ = 10^6^). Best results are seen when both constraints are applied simultaneously. Each point corresponds to a single inferred structure, and colors indicate the simulated true structure from which counts were derived. “Null” indicates inference performed without the Poisson model. Corresponding *p*-values are in Supplementary Table S1. (B) A simulated chromosome is shown alongside inferred versions of the same chromosomes using various strategies. Each panel also lists the RMSD and distance error associated with the given structure, relative to the true structure.

As an additional control for the previous experiment, we sought to confirm that the Poisson model for ambiguous diploid contact counts improved inference above what could be attained by the constraints alone. Accordingly, we compared results generated with simulated ambiguous data to “null” structures, which were inferred with the same initialization and constraints but without the Poisson model. Both measures of intra-homolog similarity showed a clear improvement when the Poisson model was incorporated in inference (Figure 2). On the other hand, the inter-homolog distance error did not improve with the addition of the Poisson model, suggesting that the constraints are the primary influence in orienting the homologs relative to one another.

### 3.2 Best results obtained by incorporation of disambiguated data

We expected that more accurate structure inference could be achieved using data where one or more ends of each contact count was disambiguated, relative to fully ambiguous data. We also expected that unambiguous data, in which both ends of each contact count are disambiguated, would yield better models than partially ambiguous data, in which only one end of each contact count is disambiguated. To test these hypotheses, we simulated partially ambiguous data and unambiguous data. Across all similarity measures, inference with unambiguous data performed best, and inference with ambiguous data performed worst, as expected (Figure 3). Partially ambiguous contacts seem especially beneficial in inference of intra-homolog structure, since intra-homolog RMSD and distance error of structures inferred with partially ambiguous counts was significantly lower than intra-homolog RMSD and distance error of structures inferred with ambiguous counts (pairwise t-test, Bonferroni corrected *p*-value <0.05, Supplementary Table S2).

**Figure 3:**
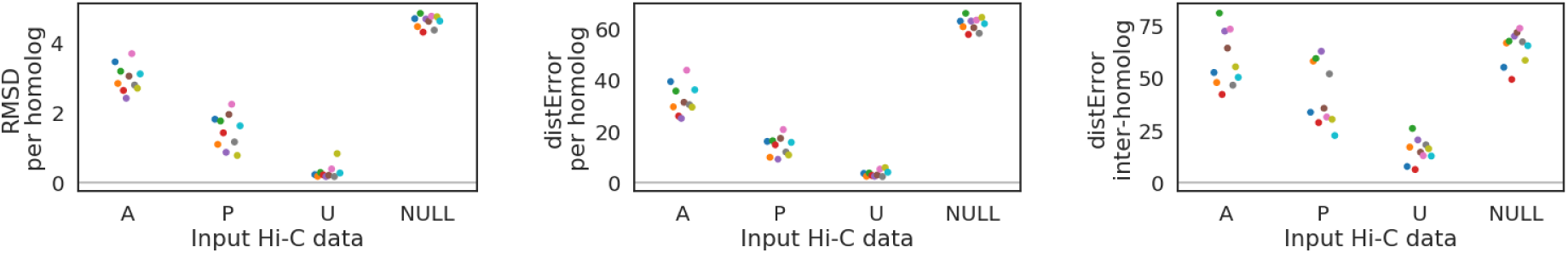
Inference with ambiguous and disambiguated data. The simulated data consists of a single chromosome with 9.3^6^ reads and 343 beads, the size of which corresponds to mouse chromosome X at 500 kb resolution. The quality of the inferred structure, as measured by three different error scores (y-axis), improves when one or both ends of a contact are disambiguated, and best results are seen in the latter case. “A” indicates ambiguous data, “U” indicates unambiguous data, and “P” indicates partially ambiguous data. Each point corresponds to a single inferred structure, and colors indicate the simulated true structure from which counts were derived. Corresponding *p*-values are in Supplementary Table S2.

### 3.3 Inference successfully identifies the superdomain structure of the inactive X chromosome

Deng et al. [10] previously showed that inactive X chromosome adopts a bipartite structure with two large superdomains, whereas the active homolog does not. We sought to validate our approach by inferring the mouse X chromosome structure and examining the degree to which each homolog exhibits a bipartite structure. Bipartite structure was assessed via the “bipartite index,” which refers to the ratio of the frequency of counts within each superdomain to those between superdomains [10]. To determine the bipartite index of an inferred 3D structure, we induced counts by applying the biophysical model used during inference (Equation 9) to the distances between beads.

We inferred a 3D structure for the mouse X chromosome at 500 kb resolution and computed the bipartite index at each bin along the chromosome. The boundary between superdomains of the inactive X chromosome has been shown to center at position 72.8–72.9 Mb (mm9, corresponding to bead 146 in our structure) [10]. In our analysis, the bipartite index of the inferred inactive homolog exhibited a prominent peak around position 75 Mb (corresponding to bead 150), whereas the active homolog only had a relatively small peak at this position (Figure 4). This observation suggests that the inference method has successfully recovered this known feature of the mouse inactive X chromosome.

**Figure 4:**
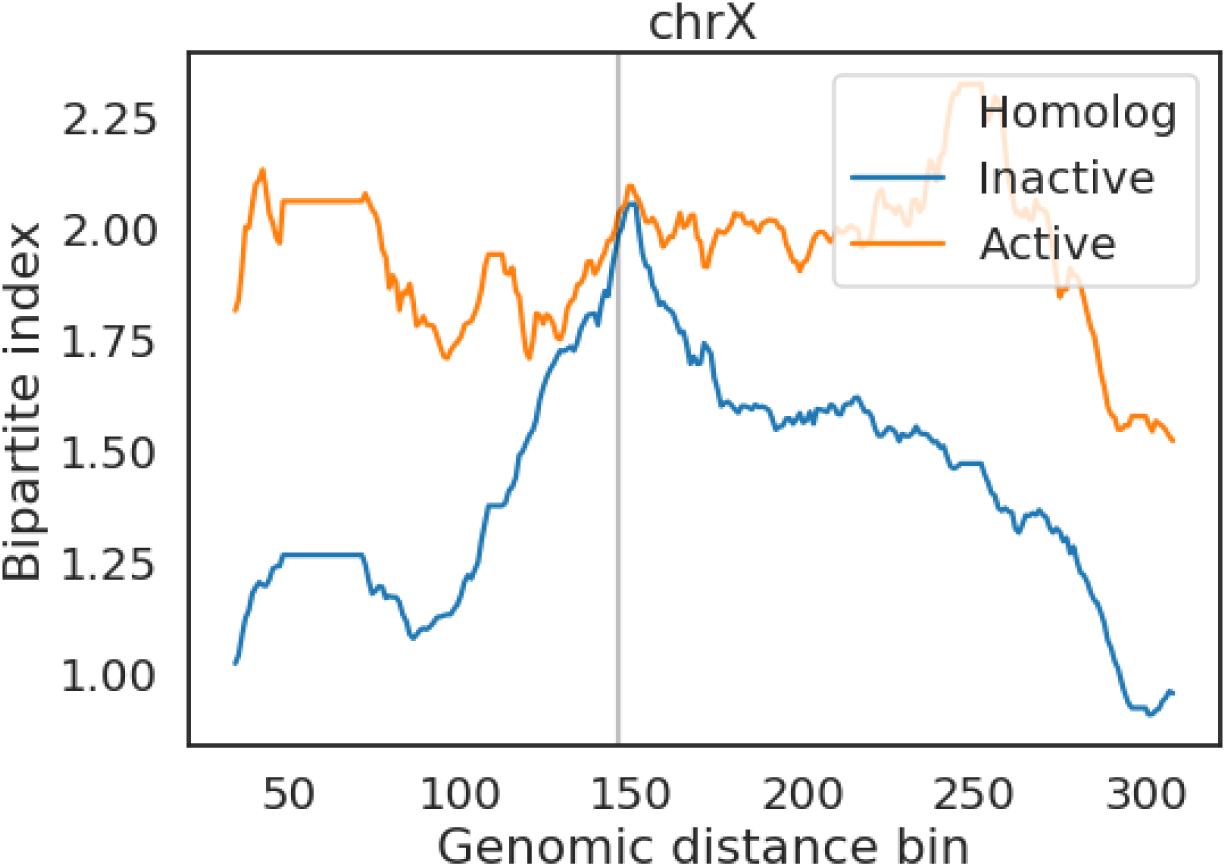
Bipartite structure of the mouse inactive X chromosome. The bipartite index (y-axis) at each genomic distance bin (x-axis) for the active (orange) and inactive (blue) homologs of the mouse X chromosome. The black line corresponds to the known boundary between superdomains of the inactive homolog at bin 146.

## 4 Discussion

Three-dimensional structural inference of diploid genomes is a challenging problem because most Hi-C data is inherently ambiguous and does not discriminate between contact counts from the two homologs of a given chromosome. Even in the rare cases when parental genotype information is available, only a minority of reads can be disambiguated. As a consequence, many inference methods have modeled a single structure per diploid chromosome [37, 22, 40]. Such an approach assumes that the two homologous copies of a given chromosome have the same 3D structure and prevents structural inference of more than one chromosome at a time.

In this work, we show how to carry out true diploid structural inference by modifying the objective function of PASTIS, a previously published haploid inference method [37]. PASTIS models each contact count via a Poisson distribution of a biophysical model between pairwise distances connecting the corresponding beads. In this work, we model each diploid contact count as the sum of biophysical models between all possible distances between the corresponding bead on each homolog. We combine this modified Poisson model with two constraints that limit the scope of possible solutions to more realistic structures. One constraint enforces even spacing of beads along the chain of the chromosome, and the other serves to spatially separate homologs. Using simulations, we show that the most accurate structures are obtained by inferring with the Poisson model in conjunction with both constraints. We note that the homologs of our simulated structures occupy distinct territories. While this is the case for many organisms, there are some exceptions [25, 32, 31, 4, 27]; therefore, the weight assigned to the homolog-separating constraint should be tuned for each organism based on prior knowledge.

A limitation to this method involves the distribution of contact count data, which may be better fit by a negative binomial model than a Poisson model [8]. Unfortunately, our method of diploid inference relies on a specific property of Poisson models, namely, that the sum of multiple Poisson variables is also a Poisson variable. Another caveat involves the biophysical model used during inference (Equation 9), which may not accurately capture the relationship between contact counts and pairwise distances in all situations. For example, this relationship may vary depending on the organism, resolution, genomic distance range, and cell cycle status [41, 1, 2, 21, 13].

We envision several ways in which diploid PASTIS could be further improved. First, diploid PASTIS could allowing for joint estimation of the *α* parameter of the biophysical model alongside the 3D structure, as is possible for haploid PASTIS. Second, results could potentially be improved by incorporating a multiscale optimization strategy, in which a high-resolution structures is inferred in a stepwise fashion through multiple rounds of inference with gradually increasing resolution. Similarly, inference of the whole genome may be improved by a stepwise approach where each chromosome is first inferred individually before being placed in the context of the whole genome.

## Acknowledgments

WSN acknowledges support from the National Institutes of Health Common Fund 4D Nucleome Program (Grant U54 DK107979). NV was supported by a BIDS fellowship from the Gordon and Betty Moore Foundation (Grant GBMF3834) and by the Alfred P. Sloan Foundation (Grant 2013-10-27).

## Supplement

**Table S1:** Constraints improve ambiguous inference. Each entry is a Bonferroni adjusted *p*-value for a *t*-test applied to the specified pair of methods. Values <0.05 are in boldface.

|  |  | RMSD<br>per homolog | Distance error<br>per homolog | Distance error,<br>inter-homolog |
| --- | --- | --- | --- | --- |
| No constraints | Bead chain connectivity | **0.0458** | **0.0104** | 0.275 |
| No constraints | Homolog separating | 1.3 | 0.222 | 4.14 |
| No constraints | Both constraints | **0.00309** | **0.00667** | **0.0179** |
| No constraints | Both constraints + Null | **0.0000404** | **0.00000773** | **0.0000883** |
| Bead chain connectivity | Homolog separating | **0.00731** | **0.00588** | 1.62 |
| Bead chain connectivity | Both constraints | 4.89 | 8.74 | 0.159 |
| Bead chain connectivity | Both constraints + Null | **0.000018** | **0.00000054** | **0.000263** |
| Homolog separating | Both constraints | **0.0018** | **0.00713** | 0.183 |
| Homolog separating | Both constraints + Null | **0.0000397** | 0.152 | 0.103 |
| Both constraints | Both constraints + Null | **0.0000179** | **0.00000387** | 0.981 |

**Table S2:** Inference with ambiguous and disambiguated data. Each entry is a Bonferroni adjusted *p*-value for a *t*-test applied to the specified pair of methods. Values <0.05 are in boldface.

|  |  | RMSD<br>per homolog | Distance error<br>per homolog | Distance error,<br>inter-homolog |
| --- | --- | --- | --- | --- |
| Ambiguous | Partially ambiguous | **0.000231** | **0.0000669** | 0.654 |
| Ambiguous | Unambiguous | **3.36E-09** | **2.88E-08** | **0.00000369** |
| Partially ambiguous | Unambiguous | **0.00000462** | **0.00000345** | **0.00000342** |

